# No evidence for angiosperm mass extinction at the Cretaceous–Paleogene (K-Pg) boundary

**DOI:** 10.1101/2023.02.15.528726

**Authors:** Jamie Thompson, Santiago Ramírez-Barahona

## Abstract

The Cretaceous-Paleogene mass extinction event (K-Pg) witnessed up to 75% of animal species going extinct, most notably among these are the non-avian dinosaurs. A major question in macroevolution is whether this extinction event influenced the rise of flowering plants (angiosperms). The fossil record suggests that the K-Pg event had a minor impact on the extinction rates of angiosperm lineages, yet the diversification of extant angiosperms was delayed and started after the K-Pg boundary. However, phylogenetic evidence for angiosperm extinction dynamics remains unexplored. Through the analyses of two angiosperm mega-phylogenies containing ~32,000–74,000 extant species, here we show relatively constant extinction rates throughout geological time and no evidence for a mass extinction at the K-Pg boundary. Despite uncertainty of earliest angiosperm branching times, their staggering diversity, and complex evolutionary dynamics, our preliminary analyses provide congruent results with the fossil record and support the macroevolutionary resilience of angiosperms to the K-Pg mass extinction.

## Introduction

Five major mass extinctions have profoundly shaped the diversity and distribution of life on Earth. The most recent of these events was the Cretaceous–Paleogene mass extinction (K-Pg) ~66 million year ago (Mya) that led to the demise of non-avian dinosaurs; as a result of the K-Pg extinction event up to 75% of vertebrate species went extinct (Raup and Sepkoski, 1982; Lyson et al., 2019) and terrestrial biomes underwent an extraordinary reconfiguration into angiosperm-dominated ecosystems (Krug et al., 2009; Carvalho et al., 2021). Today, angiosperms dominate terrestrial biomes globally, with a staggering ~290,000 species that dwarfs the ~79,000 species in the other major terrestrial plant clades (Christenhusz and Byng, 2016). The rise to ecological dominance of angiosperms accelerated after the K-Pg boundary and changed the planet and altered evolutionary trajectories of major lineages of plants, animals, and fungi (Benton et al., 2021). Angiosperms co-diversified with many lineages, including pollinating insects and herbivores (Barba-Montoya et al., 2018), but their success drove extinction in others, including the conifers (Condamine et al., 2020).

Fossilised stem relatives of angiosperms are known from deposits ranging from the Permian to the Late Cretaceous, but the age of origin of crown angiosperms remains uncertain (Sauquet et al., 2022). However, the fossil record and phylogenetics provide robust evidence on the later history of the group, particularly since the Early Cretaceous. Most extant angiosperm lineages originated in the Cretaceous, but whether their extinction dynamics were altered by the K-Pg mass extinction remains unknown. While fossils reveal the demise of non-avian dinosaurs at K-Pg and high extinction rates in other lineages including birds (Longrich et al., 2011) and reptiles (Longrich et al., 2012), different trajectories are observed in major plant lineages. High extinction rates follow the K-Pg event in non-flowering seed plants, and before this event in spore-bearing plants (Silvestro et al., 2015). In contrast, the angiosperm fossil record indicates relatively low and stable extinction rates through time and no evidence of shifting rates across the K-Pg boundary.

Fossils provide a physical record of evolutionary dynamics, but for angiosperms it is sparse and fossils can only be assigned to ~257 of ~13,164 extant angiosperm genera (Silvestro et al., 2015). Additionally, the fossil record has many sources of bias through time and space (Xing et al., 2016). Thus, whether angiosperm diversity was shaped by the K-Pg event remains as one of six major unanswered questions in flowering plant evolution (Sauquet and Magallón, 2018). Angiosperm extinction around the K-Pg event are hard to gauge with fossils alone and the story remains incomplete without phylogenetic evidence, which otherwise would provide key insights into the evolution of modern ecosystems worldwide.

By applying phylogenetic comparative methods to molecular sequences we can detect signatures of mass extinctions in large phylogenies of extant species (May et al., 2016; Kopperud et al., 2023) and gain macroevolutionary insights in the face of substantial taphonomic biases (lineages and regions with a poor fossil record; Hernández-Hernández et al., 2014). However, phylogenies are reconstructions, not observations, and suffer from multiple sources of uncertainty (Rangel et al., 2015). This is especially true for angiosperm due to their extreme richness, complex evolution, and uncertain age (Sauquet et al., 2022). A major source of uncertainty concerns the backbone topology of angiosperms and the timings of early divergences, both of which remain uncertain (Sauquet and Magallón 2018). Estimates of the crown group age of angiosperms range from 140 to 270 million years and support for both extremities come from fossil and molecular data (for details, see Sauquet et al., 2022). Accounting for these sources of uncertainty by analysing a sample of plausible phylogenies is currently intractable on the scale of all angiosperms, but available mega-phylogenies capture wide variation in age estimates while comprehensively sampling extant diversity.

Even though phylogenetic data and methods are available to assess mass extinctions in angiosperms, these question remains poorly unexplored (Silvestro et al., 2015; Sauquet and Magallón, 2018). Here, we applied the Bayesian method CoMET (May et al., 2016) to estimate mass extinctions and assess the influence of the K-Pg event on angiosperm extinction rates. To capture some of the phylogenetic and age uncertainties we analysed the two angiosperm-wide mega-phylogenies produced by Zanne et al. (2013) and Smith and Brown (2018), which differ both in relationships and divergence ages. Both mega-phylogenies sample a very large portion of angiosperm diversity: ~32,000 species (~10.5%) in Zanne et al. (2013) and over 70,000 (~25.2%) in Smith and Brown (2018). Backbone topology differs somewhat among these trees, but estimated ages of the earliest branches differs considerably, which has knock-on effects for more recent divergence timings. Zanne et al. (2013) time-calibrated the angiosperm phylogeny based on Soltis et al. (2011) and estimated a crown age of ~243.3 Mya, whereas Smith and Brown (2018) time-calibrated their phylogeny based on Magallón et al. (2015) to estimate a crown age of ~139.4 Mya. These timings span the plausible interval of 140–270 Mya for crown angiosperms (Sauquet et al., 2022) and provide two distinct but credible evolutionary histories for analyses into impacts of the K-Pg extinction event (Figure 1).

**Figure 1.**
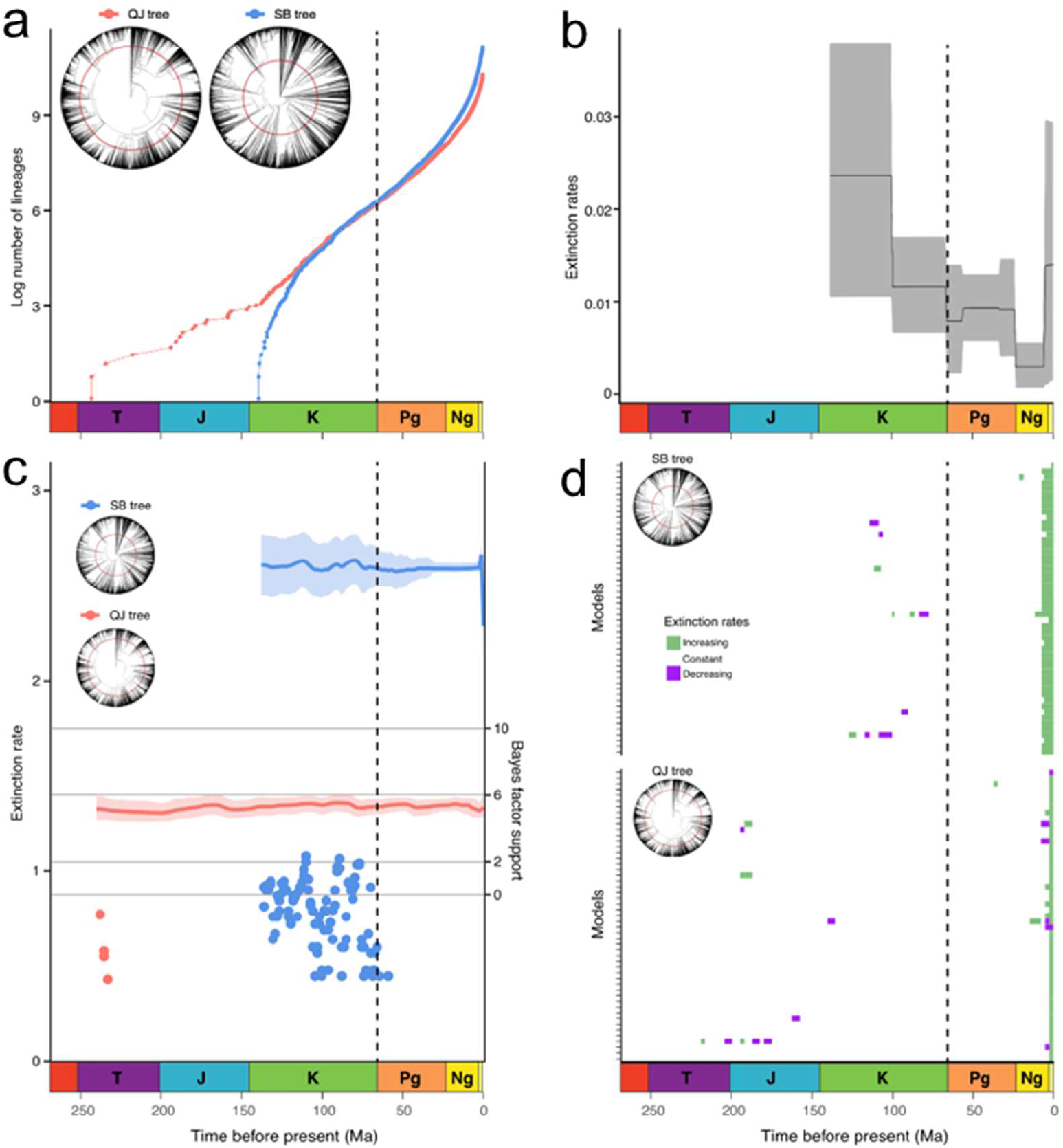
Macroevolutionary dynamics of angiosperms not negatively impacted at the K-Pg mass extinction event. Lineage through time plots (a) and phylogenies comprising ~32,000 and ~74,000 species (denoted as QJ and SB, respectively). Generic-level extinction rates of angiosperms estimated with PyRate (b), reproduced with permission from Silvestro et al. (2015). Phylogenetic extinction rates estimated by CoMET, with detected mass extinctions (c), none of which are strongly supported (Bayes factor >6). Trends in alternative models in congruence classes of both analyses (d). The geological timescale is visualised in each panel, and K-Pg is represented by vertical dashed lines in the plots and concentric lines in the phylogenies.

## Results and discussion

Our Bayesian analyses revealed relatively stable extinction rates through time and no evidence of mass extinctions at any time during the last 150 million years (Figure 1). Furthermore, we find overwhelming Bayes factor (BF) support for constant diversification models over mass extinctions (SB phylogeny BF >65,000; QJ BF > 268,000). This agrees with fossil evidence of angiosperms suggesting relatively stable extinction rates and no impacts of the K-Pg mass extinction on flowering plant diversity (Silvestro et al., 2015), despite significant reconfigurations of terrestrial habitats into the Paleogene (Carvalho et al., 2021). In fact, the K-Pg extinction event may have played a role in favouring the emergence of angiosperm-dominated ecosystems; fossil evidence links the K-Pg to the rise of the modern Amazonian rain forests (Carvalho et al., 2021).

The staggering extant biodiversity of angiosperms was not shaped by variable extinction rates (Figure 1, Silvestro et al., 2015). Instead, the rise of angiosperms is likely better explained through numerous radiations driven by the acquisition of eco-morphological innovations and associated with climate change and ecological opportunity (Soltis et al., 2019; Eriksson et a. 2000; Friis et al. 2006). Diverse reproductive strategies, including wind- and insect-pollination, enabled survival in changing environments (McElwain and Punyasena, 2007) and variation in seed dispersal modes (e.g., wind, water, and animals) facilitated the colonisation of newly opened areas. Additionally, the wide range of ecological niches (tropical to arid environments) inhabited by angiosperms since the Late Cretaceous (Ramírez-Barahona et al., 2020; Benton et al., 2021) promoted lineage diversification prior to the K-Pg boundary and the eventual survival of angiosperm lineages when confronted with major environmental changes associated with the K-Pg.

Our preliminary analyses evince the resilience of angiosperms to the K-Pg extinction event, contrasting with phylogenetic evidence for other lineages such as nonavian dinosaurs, conifers (May et al., 2016), and fishes (Arcila and Tyler., 2017). Although phylogenies provide data for >90% of angiosperm lineages that did not fossilise, these trees are not observations and phylogenetic reconstructions are sensitive to numerous sources of uncertainty. For instance, different reconstructions for mammals support contrasting models for extinction rates across the K-Pg boundary. A nearly complete tree of extant mammals suggests stability of diversification rates across K-Pg (Bininda-Edmonds et al., 2007), whereas a molecular phylogeny calibrated with a relaxed clock indicates important roles of the K-Pg in shaping macroevolutionary trends (Meredith et al., 2011). Analyses of diversification in deep time suffer from nonidentifiability (Louca and Pennell., 2020), in which infinite combinations of speciation and extinction rates can fit the same phylogeny with the same likelihood scores (congruent classes). To assess the impact of non-identifiability and assess shared features among models within the same congruence class (Höhna et al., 2022), we explored congruence classes for the two mega-phylogenies and found agreement across multiple models in both phylogenies, which supported constant extinction trends across the K-Pg boundary (Figure 1).

The uncertainty of earliest angiosperm branching times, their staggering diversity, and complex evolutionary dynamics present major challenges to accounting for uncertainty when exploring deep-time dynamics. Nonetheless, our preliminary analyses provide congruent results between two very different mega-phylogenies and strengthens the argument of the macroevolutionary resilience of angiosperms to the K-Pg mass extinction as revealed by fossils.

The K-Pg extinction event caused widespread plant extinction and changes in ecosystem composition at local and regional scales, such as tropical rain forests (Carvalho et al., 2021), and recovery from this disruption varied across Earth and lineages (Stiles et al., 2020). While angiosperms as a whole appeared to have had resilient macroevolutionary dynamics to the K-Pg event relative to other plant and animal lineages, it is important to understand the heterogeneous impacts this event may have had in different angiosperm lineages. Applying the methods used here to particular angiosperm lineages could reveal variation in macroevolutionary resilience associated with geographic ranges and eco-morphological traits, such as high-latitudinal distributions (McLoughlin et al., 2008), insect-pollination (McElwain and Punyasena, 2007), and polyploidy (Fawcett et al., 2009).

Characterising the impact of past extinctions on lineage dynamics is vital to predict how biodiversity will fare under current anthropogenic ecosystem changes (Pyron and Pennell, 2022), which is especially important in angiosperms, upon which human existence depends (Tilman et al., 2002). Whether lineages characterised by particular geographic ranges and eco-morphological traits show different macroevolutionary dynamics to those without will not only inform longstanding macroevolutionary debates, but could help direct conservation efforts to protect vulnerable lineages in the face of the hypothesised sixth mass extinction (Ceballos and Ehrlich, 2018).

## Material and methods

### Diversification analyses

We used updated versions of the molecular mega phylogenies produced by Zanne et al. (2013) and Smith and Brown (2018); non-angiosperm species were pruned prior to analyses. Qian and Jin (2016) ccorrected the taxonomy of the Zanne et al. (2013) mega phylogeny, removed duplicates, and added six families to extend coverage to all currently recognized families. Igea and Tanentzap (2020) standardised the taxonomy of the Smith and Brown (2018) mega phylogeny against ‘The Plant List V1.1’ (Jin and Qian, 2019).

We produced log-transformed lineage through time (LTT) plots for both megaphylogenies using the *phytools* package (Revell 2012) in R (R Core Team 2023). We compared support for models of constant diversification and mass extinction at ~66 Mya in both phylogenies with marginal likelihoods, estimated with stepping-stone sampling in the TESS package in R (Höhna et al., 2016). In each model we specified the fraction of sampled species, ran 1,000 iterations with burn-in of 100, implemented 50 steppingstones, and estimated Bayes factor support between models.

We estimated diversification dynamics with mass extinctions using the CoMET model (May et al., 2016) implemented in TESS. A threshold of instantaneous species-loss of 75% with a beta-distribution for survival probability spanning ~18-~32% was implemented using a compound Poisson process; a loss of 75% diversity is a relatively relaxed threshold but agrees with estimates of species loss during the K-Pg event. Incomplete sampling was accounted for by specifying the fraction of sampled species, the number of expected rate changes was set to 25 (slightly more conservative than Magallón et al., 2019), the number of expected mass extinctions was set to one, and empirical hyperpriors for speciation and extinction rates were estimated automatically with an initial MCMC run. The final analyses were run in replicate to convergence until an effective sample size >300 was achieved; the first 10,000 generations of each run were discarded as burn-in.

### Sensitivity testing

Sensitivity testing was performed in CoMET using the smaller Zanne et al. (2013) phylogeny by replicating analyses and altering the number of expected rate changes five times from 1–1000 (1,100,200,500,1000). Further analyses were undertaken to account for the non-identifiability of diversification rates (Louca and Pennell 2020) with the CRABS (Höhna et al., 2022) package in R. Congruence classes of the CoMET parameters were explored with 500 models, assuming a constant rate of extinction, and in the absence of any linear or exponential temporal trend. We sampled and visualised 50 models for each mega phylogeny to reduce crowding in the figure.

